# Molecular Architecture of Early Dissemination and Evolution of the SARS-CoV-2 Virus in Metropolitan Houston, Texas

**DOI:** 10.1101/2020.05.01.072652

**Authors:** S. Wesley Long, Randall J. Olsen, Paul A. Christensen, David W. Bernard, James R. Davis, Maulik Shukla, Marcus Nguyen, Matthew Ojeda Saavedra, Concepcion C. Cantu, Prasanti Yerramilli, Layne Pruitt, Sishir Subedi, Heather Hendrickson, Ghazaleh Eskandari, Muthiah Kumaraswami, Jason S. McLellan, James M. Musser

## Abstract

We sequenced the genomes of 320 SARS-CoV-2 strains from COVID-19 patients in metropolitan Houston, Texas, an ethnically diverse region with seven million residents. These genomes were from the viruses causing infections in the earliest recognized phase of the pandemic affecting Houston. Substantial viral genomic diversity was identified, which we interpret to mean that the virus was introduced into Houston many times independently by individuals who had traveled from different parts of the country and the world. The majority of viruses are apparent progeny of strains derived from Europe and Asia. We found no significant evidence of more virulent viral types, stressing the linkage between severe disease, underlying medical conditions, and perhaps host genetics. We discovered a signal of selection acting on the spike protein, the primary target of massive vaccine efforts worldwide. The data provide a critical resource for assessing virus evolution, the origin of new outbreaks, and the effect of host immune response.

**Significance:** COVID-19, the disease caused by the SARS-CoV-2 virus, is a global pandemic. To better understand the first phase of virus spread in metropolitan Houston, Texas, we sequenced the genomes of 320 SARS-CoV-2 strains recovered from COVID-19 patients early in the Houston viral arc. We identified no evidence that a particular strain or its progeny causes more severe disease, underscoring the connection between severe disease, underlying health conditions, and host genetics. Some amino acid replacements in the spike protein suggest positive immune selection is at work in shaping variation in this protein. Our analysis traces the early molecular architecture of SARS-CoV-2 in Houston, and will help us to understand the origin and trajectory of future infection spikes.

## Introduction

Pandemic disease caused by the severe acute respiratory syndrome coronavirus 2 (SARS-CoV-2) virus is now responsible for massive human morbidity and mortality worldwide (1–4). The virus was first documented to cause severe respiratory infections in Wuhan, China, beginning in late December 2019 (5–7). Global dissemination occurred extremely rapidly, and has affected major population centers on many continents, especially in Asia, Europe, and North America (8–10). In the United States, Seattle and the New York City region have been especially important centers of COVID-19 disease caused by SARS-CoV-2. For example, as of April 15, 2020, there were 111,424 confirmed SARS-CoV-2 cases in NYC, causing 29,741 hospitalizations and 10,899 fatalities (11). Similarly, in Seattle and King County, 4,620 positive patients and 303 deaths have been reported as of April 15, 2020 (12).

The Houston metropolitan area is the fourth largest and the most ethnically diverse city in the United States, with a population of approximately 7 million (13). The 2,000-bed Houston Methodist Health System has eight hospitals and serves a large multinational and socioeconomically diverse patient population throughout greater Houston. Although the city is well-known as the energy capital of the world, Houston also has a very large port, and the George Bush International Airport is a major transportation hub for domestic and international flights to Asia, Europe, and Central and South America. Many of these flights provide direct city-to-city service to diverse countries, including global population centers. The first COVID-19 case in metropolitan Houston was reported on March 5, 2020 (14). Community spread was suspected of occurring one week later (14). Many of the first cases in our region were associated with national or international travel, including areas known to have COVID-19 virus outbreaks (14). These facts, coupled with the existence of a central molecular diagnostic laboratory that serves all Houston Methodist hospitals and our very early adoption of a molecular test for the SARS-CoV-2 virus, permitted us to rapidly interrogate genomic variation among strains causing infections in the greater Houston area.

We here report that SARS-CoV-2 was introduced to the Houston metropolitan region many independent times from diverse geographic regions, including Europe, Asia, and South America. The virus spread rapidly and caused disease throughout the metropolitan region. We identified clear genomic signals of person-to-person transmission. Some events were known or inferred based on conventional public health maneuvers, but many were not. In addition, spatial-temporal mapping found evidence of rapid and widespread community dissemination soon after COVID-19 cases were reported in Houston. Analysis of the relationship between distinct genomic viral clades and hospitalization did not reveal significant evidence of more virulent genome types, underscoring the need for a better understanding of the relationship between severe disease, underlying medical conditions, gender, and host genetics.

## Results

### Description of metropolitan Houston

Houston, Texas, is located in the southwestern United States, 50 miles inland from the Gulf of Mexico. It is the most ethnically diverse city in the United States (15). Metropolitan Houston is comprised predominantly of Harris County plus parts of several contiguous surrounding counties (Austin, Brazoria, Chambers, Fort Bend, Galveston, Liberty, Montgomery, and Waller). In the aggregate, the metropolitan area includes 9,444 sq. mi. The estimated population size of metropolitan Houston is 7 million (13).

### Epidemiologic curve

The first confirmed case of COVID-19 in the Houston metropolitan region was reported on March 5, 2020 (14), and the first confirmed case diagnosed in the Houston Methodist Hospital System was reported on March 6, 2020. Through April 15, 5,602 COVID-19 cases have been reported in Houston, including 1,097 cases in the Houston Methodist Hospital System (**Fig. 1**). During the study period (March 5 through March 30, 2020), the Molecular Diagnostic Laboratory in the Department of Pathology and Genomic Medicine tested 3,080 specimens, of which 406 were positive (13.2%), representing 40% (358/898) of all confirmed cases in metropolitan Houston. This large percentage is likely due to the fact that the Houston Methodist Hospital laboratory was the first hospital-based facility to have molecular testing capacity available.

**Fig. 1.**
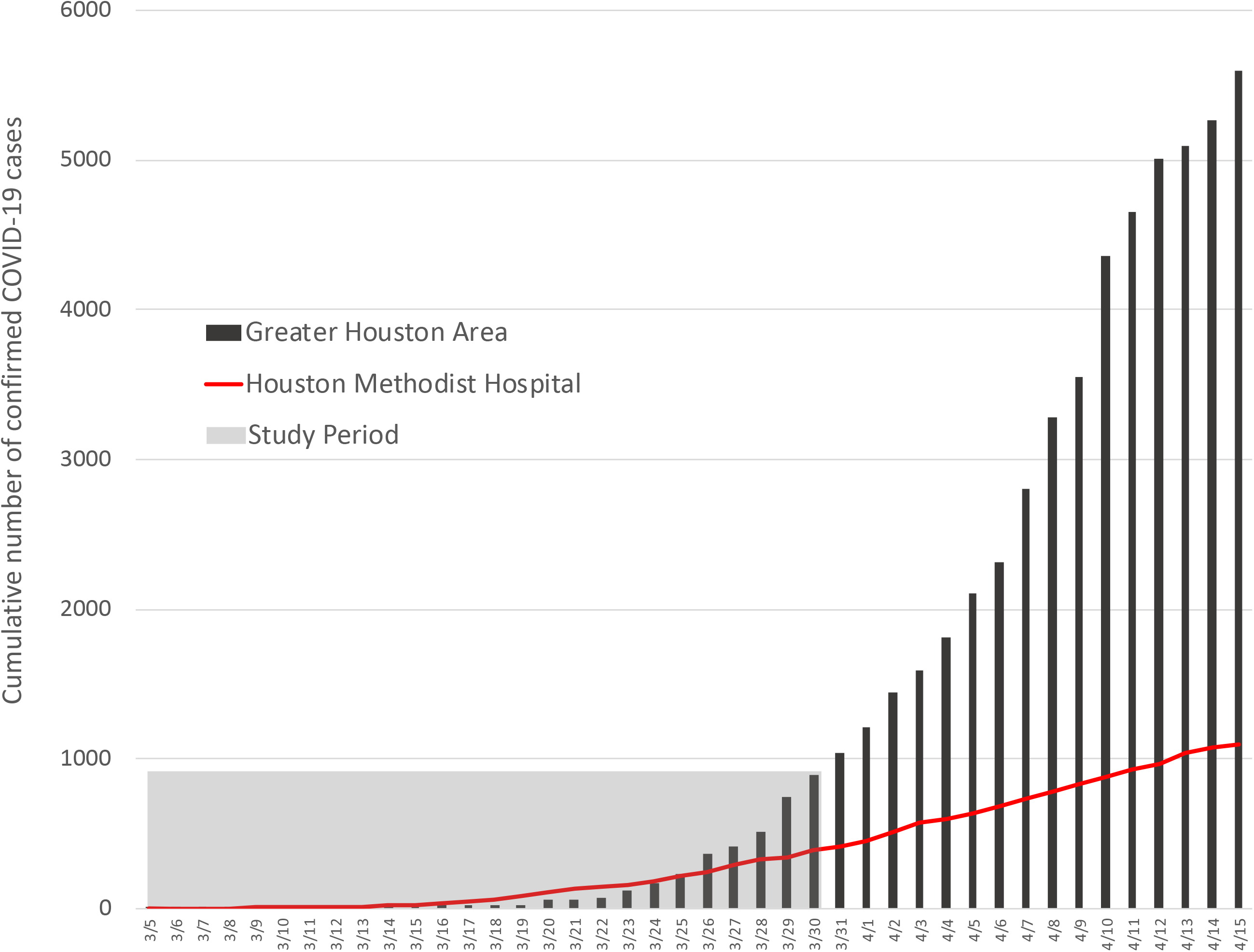
Infection rate in the Greater Houston Metropolitan region. Number of COVID-19 patients over time through April 15, 2020. Counties include Austin, Brazoria, Chambers, Fort Bend, Galveston, Harris, Liberty, Montgomery, and Waller. The shaded area represents the time period during which viral genomes characterized in this study were recovered from COVID-19 patients. The red line represents the number of COVID-19 patients diagnosed in the Houston Methodist Hospital Molecular Diagnostic Laboratory.

### Viral genome sequencing and phylogenetic analysis

We sequenced the genomes of SARS-CoV-2 strains dating to the earliest stages of confirmed COVID-19 disease in Houston. Phylogenetic analysis identified the presence of many diverse viral genomes that in the aggregate represent many of the major clades identified to date (16) (**Fig. 2**). Clades A2a, B, and B1 were the three more abundantly represented phylogenetic groups (**Fig. 2**).

**Fig. 2.**
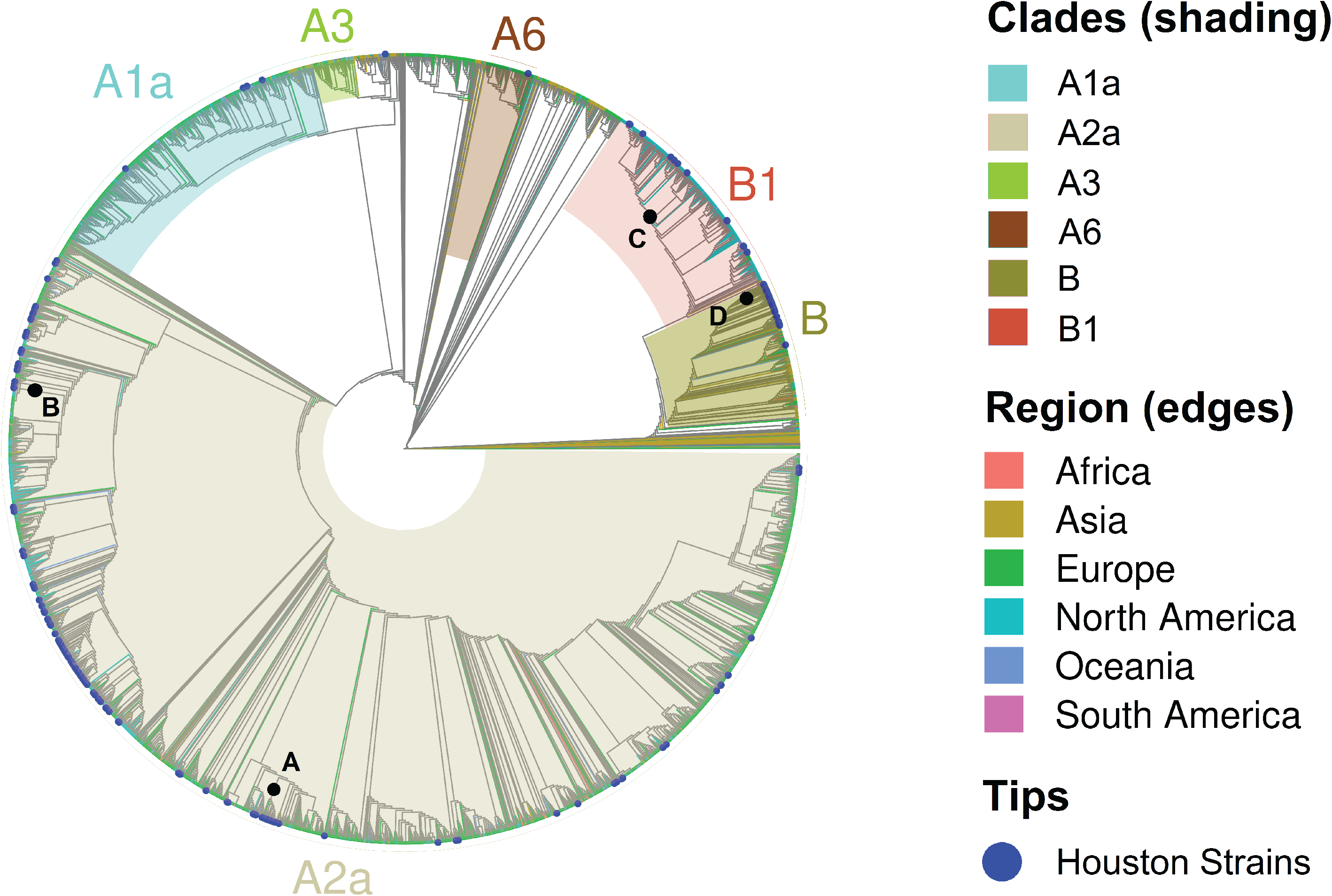
Phylogenetic relationships among Houston Methodist Hospital SARS-CoV-2 strains and relationship with strains from global sources. Phylogenetic relationships among Houston Methodist Hospital SARS-CoV-2 genome sequences and those deposited in GISAID as of April 13, 2020. Clades A1a, A2a, A3, A6, B, and B1 are color-shaded. The edge branches are colored based on region of patient origin listed in GISAID, that is, the geographic location where the sample was collected. The blue dots at the tips represent metropolitan Houston strains characterized in this study. The black dots (labeled A-D) represent the root branches that define each subclade represented in Fig. 4.

### Geospatial and time series

We examined the spatial and temporal mapping of the genome data to investigate community spread (**Fig. 3**). The figure illustrates evidence of widespread and rapid community dissemination soon after the initial COVID-19 cases were reported in Houston. There is also evidence to suggest there were multiple independent strains introduced into metropolitan Houston, followed by local spread throughout all regions of the community.

**Fig. 3.**
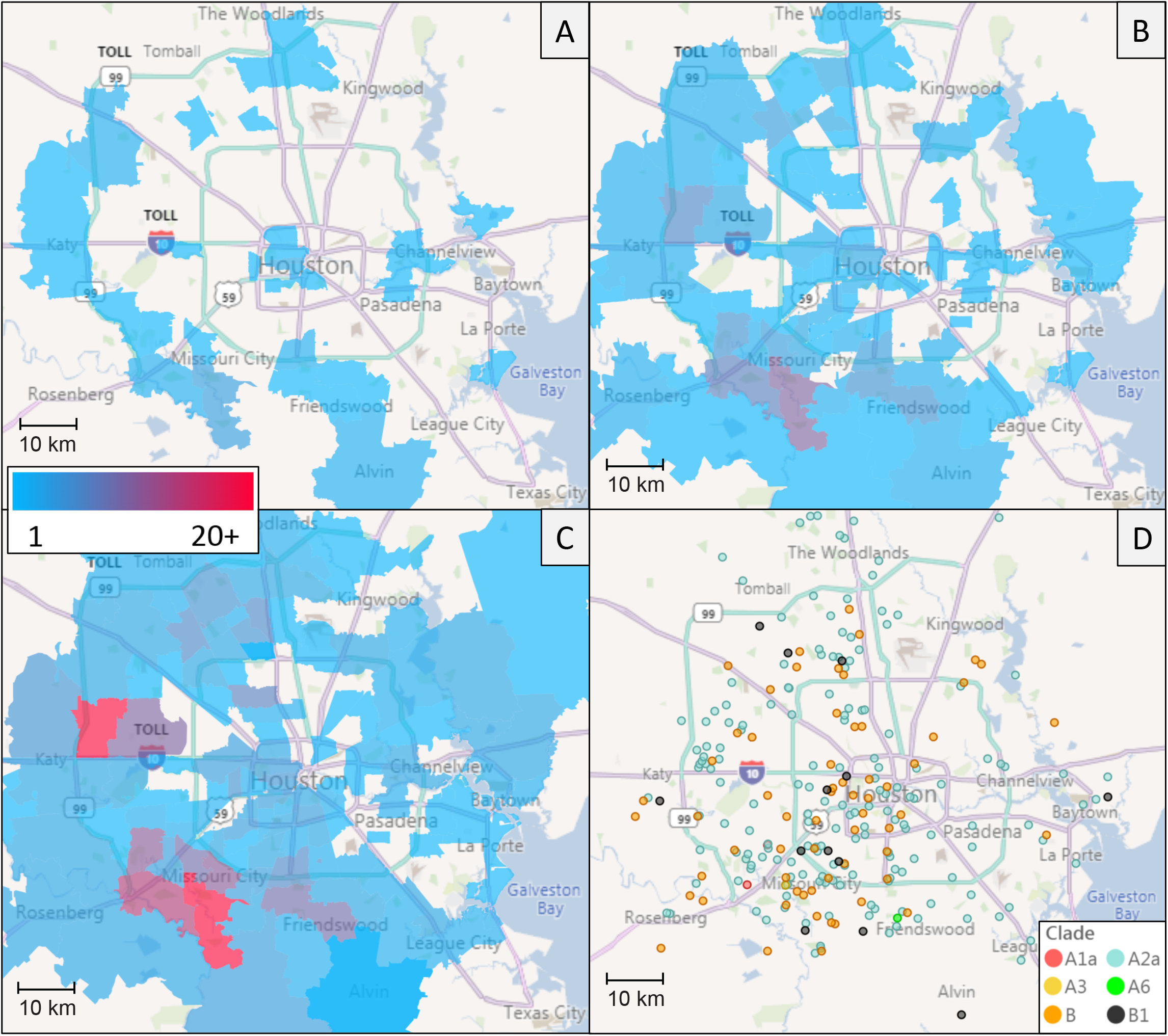
Sequential time-series heatmaps for all COVID-19 Houston Methodist patients during the study period (A-C). Geospatial distribution of sequenced SARS-CoV-2 strains (D). (*A*) Collection dates between 3/5/2020 - 3/16/2020. (*B*) Collection dates between 3/5/2020 - 3/23/2020. (*C*) Collection dates between 3/5/2020 - 3/30/2020. (*D*) Geospatial distribution of COVID-19 patients based on zip code, with colors representing the sequenced viral clade.

### Epidemiologically linked patients

We investigated the relationships among some of the genomes that were obtained from patients known to share common epidemiologic associations, such as living in the same household. In all instances, individuals known to be epidemiologically associated had identical or nearly-identical SARS-CoV-2 genomes, a finding consistent with person-to-person transmission of the virus.

### Geospatial relationships of genetically similar isolates

We next tested the hypothesis that genetically related viruses were constrained to particular geographic areas of metropolitan Houston. Although in some instances this was the case, an important observation was that most of the individual related subclades were comprised of strains distributed over broad geographic areas (**Fig. 4**). These findings are consistent with the known propensity of respiratory virus SARS-CoV-2 to spread rapidly from person to person. Patients with shared viral genomes likely constitute epidemiologic clusters, as a consequence of direct transmission to one another, shared transmission through an unknown third party, or via a reticulate network.

**Fig. 4.**
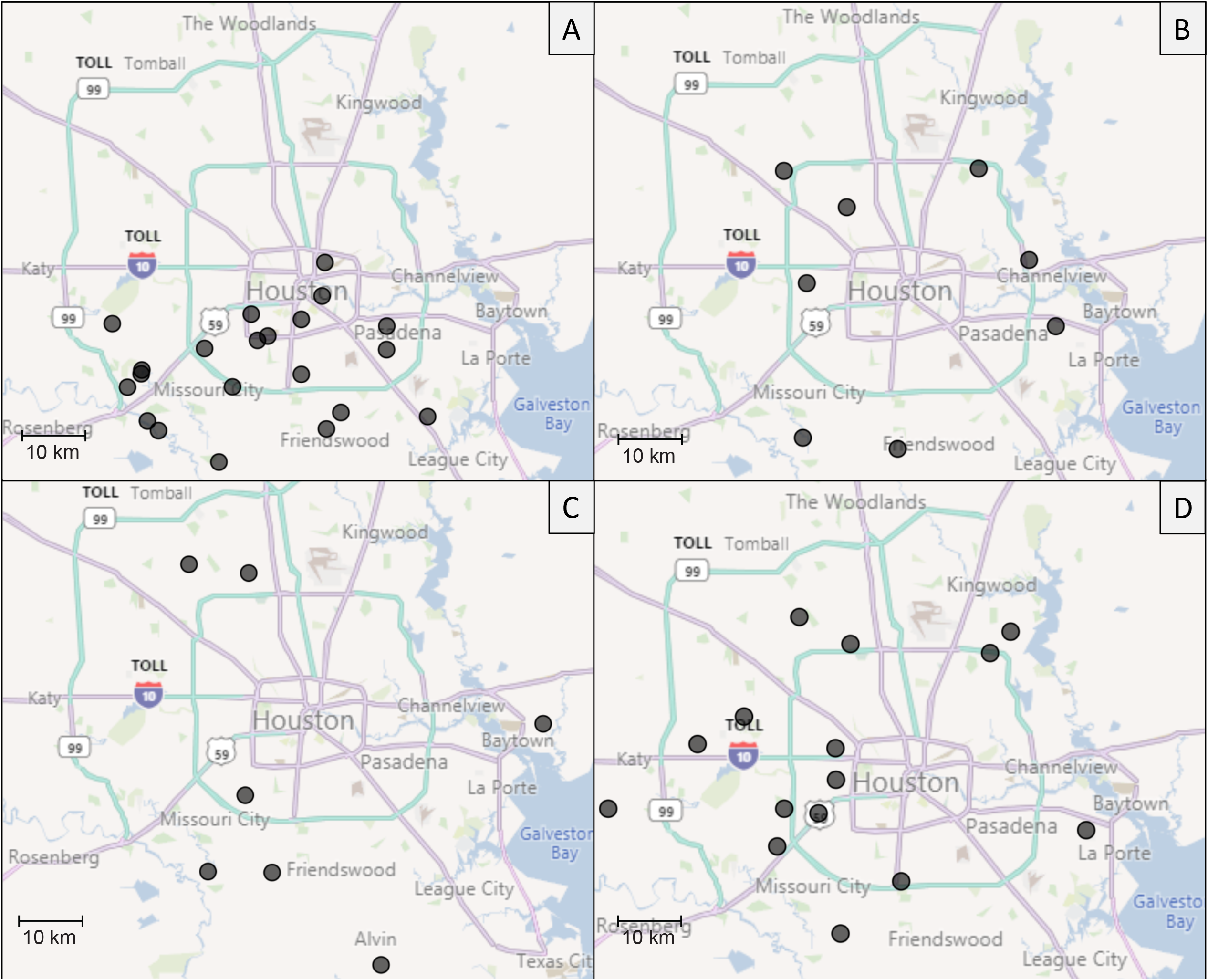
Geospatial relationship of genetically similar isolates. Each panel (A-D) illustrates a subclade of genetically similar isolates mapped against the location of each patient (black dots). The phylogenetic root branch for each subclade is labeled on the tree diagram shown in Fig. 2.

### Relationship between viral clades, clinical characteristics of infected patients, and other metadata

It is possible that SARS-CoV-2 genetic subtypes may have different clinical characteristics, analogous to what was thought to have occurred with the Ebola virus (17–19) and is known for other infectious agents. As an initial examination of this issue in SARS-CoV-2, we tested the hypothesis that patients with disease severe enough to warrant hospitalization were infected with a non-random subset of virus genotypes. We also tested the hypothesis of non-random association between virus clades and disease severity based on in-patient versus out-patient status, the need for mechanical ventilation, and the number of days on a ventilator. We found no apparent simple relationship between viral clades and disease severity using these indicators of disease severity. Similarly, there was no simple relationship between viral clades and other metadata, such as gender, age, or length of stay (**Fig. S1-S7**).

### Machine learning analysis

Machine learning models were built to predict mortality, length of stay, in-patient status, or ICU admission based on the viral genome sequence. F1 scores (the evaluation metric used in classification algorithms) for most classifiers ranged between 0.5 to 0.6, indicative of classifiers that are performing similarly to random chance. Outcome (lived versus died) was only weakly correlated with age in this data set (PCC = 0.318), and similarly regression models built to predict age and length of stay based on the viral genome sequence also had poor performance, with R^2^ values near zero. Classifiers were also trained to predict host characteristic, gender and ethnic group. The two largest ethnic groups in the patient population were African American and Caucasian, as recorded in the electronic medical record. The binary classification model had an F1 score of 0.67 [0.51-0.83, 95% CI], likely reflecting social networks in early person-to-person transmission. A table of classifier accuracy scores and performance information is provided as **Table S1**.

### Analysis of the nsp12 polymerase gene

The SARS-CoV-2 genome encodes an RNA dependent RNA polymerase (RdRp) used to replicate this RNA virus. Two amino acid substitutions (Phe476Leu and Val553Leu) in the nsp12 RdRp have been reported to confer significant resistance *in vitro* to remdesivir, an adenosine analog (20). Remdesivir is inserted into RNA chains by RdRp during replication, resulting in premature termination and inhibition of virus. This compound has shown prophylactic and therapeutic benefit against MERS-CoV experimental infection in rhesus macaques (21). A recent publication describing results from a compassionate use protocol reported that remdesivir may have therapeutic benefit in some hospitalized patients (22). If efficacy is confirmed by a randomized controlled trial, this drug may be be widely used in large numbers of patients worldwide.

To acquire baseline data about allelic variation in the gene encoding nsp12 RdRp, we analyzed the 320 sequenced viral genomes. The analysis identified 12 nonsynonymous single nucleotide polymorphisms (SNPs) in nsp12, resulting in 11 amino acid replacements throughout the protein (**Table 1**). The most common amino acid change was Pro322Leu, identified in 224 of the 320 Houston isolates. This amino acid replacement is present in genomes from the A2a clade, and distinguishes the A2a clade from other SARS-CoV-2 clades. The other amino acid changes in nsp12 were mainly present in single isolates from individual Houston strains, and some have been identified in other strains in the global GISAID collection (23, 24).A prominent exception was the identification of 29 strains with a Met600Ile polymorphism. All 29 strains were phylogenetically very closely related members of the A2a clade, and they all also had the P322L amino acid replacement characteristic of this clade (**Fig. S8**). These data indicate that the Met600Ile change is the derived state among strains with the Pro322Leu replacement. Importantly, none of the observed amino acid polymorphisms in nsp12 were located precisely at the two sites associated with remdesivir *in vitro* resistance (20), and 10 of the 11 polymorphic amino acids are located on the surface of this RdRp. However, importantly, the Ala448Val replacement occurs in an amino acid that is located directly above the nucleotide substrate binding site that is comprised of Lys545, Arg553, and Arg555, as recently shown by structural studies (25, 26) (**Fig. 5**). The Ala448 position is comparable to Val553 (**Fig. 5**), and a Val553Leu mutation in SARS-CoV was identified to confer resistance to remdesivir (20).

**Table 1.**
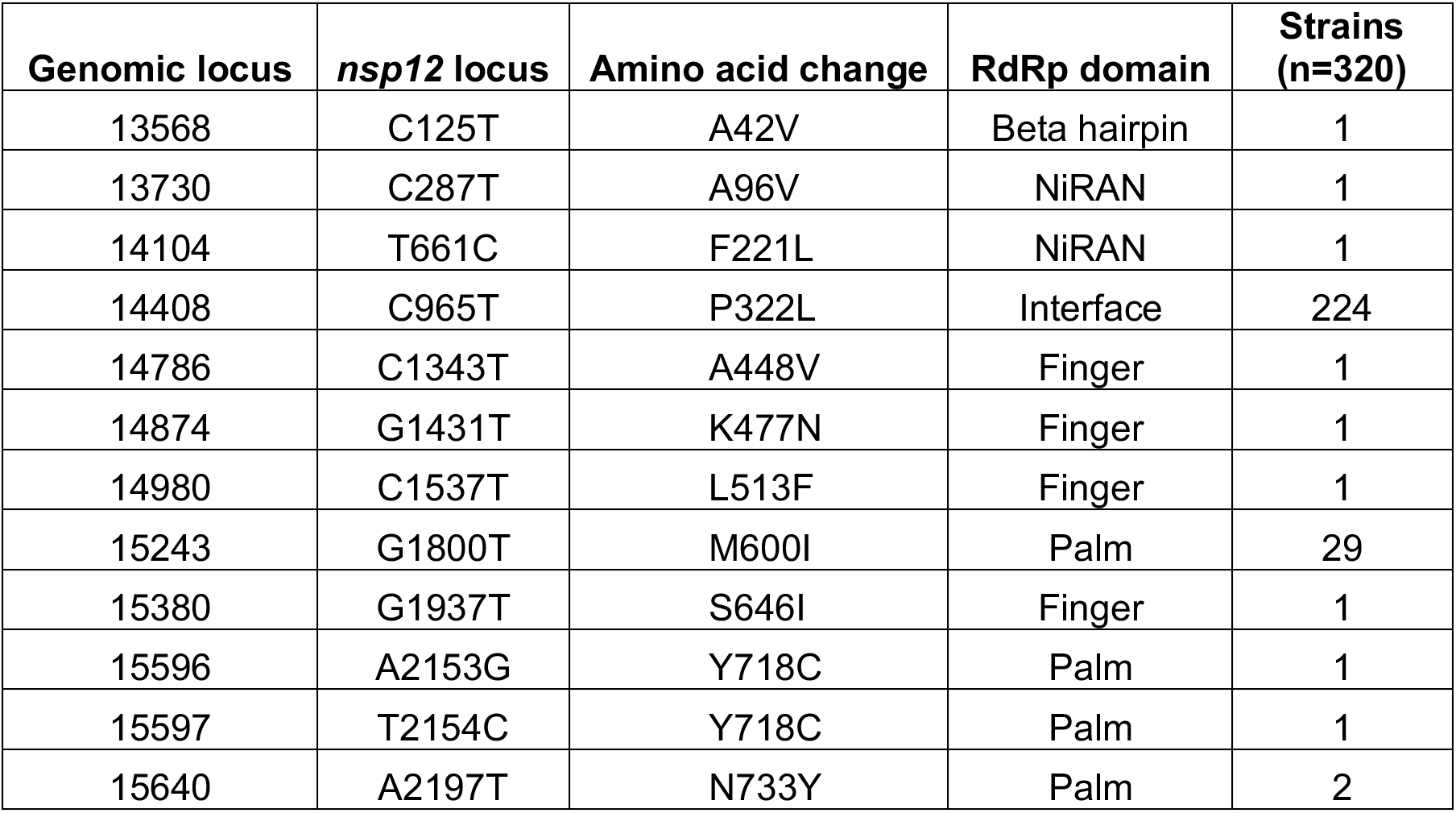
Nonsynonymous SNPs of SARS-CoV-2 *nsp12*.

**Fig. 5.**
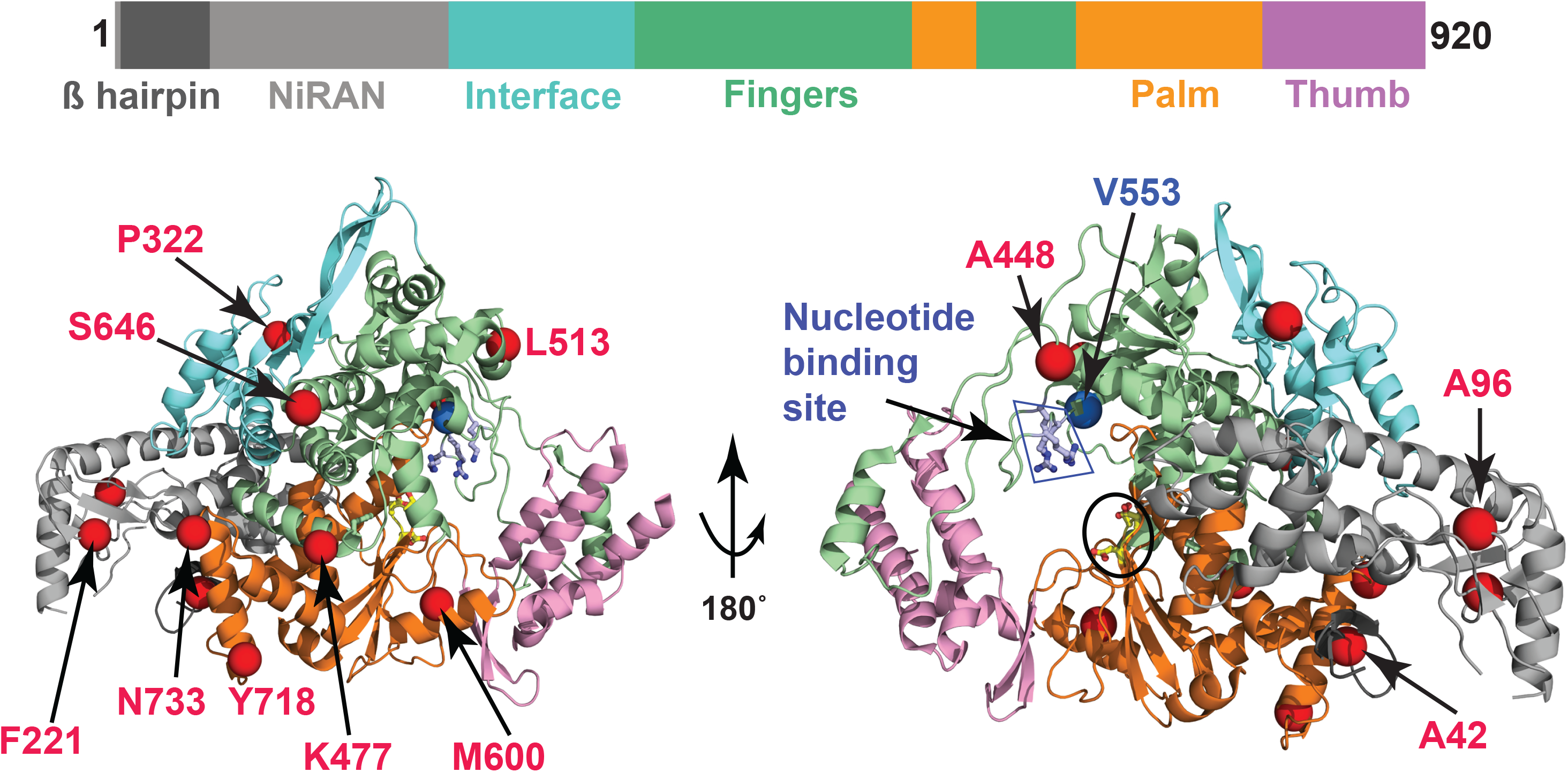
Polymorphic amino acids identified in nsp12 in this study. Nsp12 is the target for remdesivir, a nucleoside analog that may have therapeutic benefit for COVID-19 patients. The schematic shows the domain architecture of nsp12. The individual domains of nsp12 are color-coded and labeled. Ribbon representation of the crystal structure of nsp12 is shown (PDB code: 6M71). The structure in the right panel is obtained by rotating the left panel 180° along the y-axis. The nsp12 domains are colored as in the schematic at the top. The positions of Cα atoms of the 11 variant amino acids identified in this study are shown as red spheres and labeled. The catalytic site in RdRp is marked by a black circle in the right panel. The side chains of amino acids comprising the catalytic site of RdRp are shown as balls and sticks and colored yellow. The nucleotide binding site is boxed and labeled in the right panel. The side chains of amino acids participating in nucleotide binding (Lys545, Val553, and Arg555) are shown as balls and sticks. The location of Cα atoms of remdesivir resistance-conferring amino acid V553 is shown as a blue sphere and labeled.

### Analysis of the gene encoding the spike protein

SARS-CoV-2 and related coronavirus such as SARS-CoV and MERS gain entry into susceptible host cells using a densely glycosylated viral-surface molecule knows as spike (S) protein. The S protein of SARS-CoV-2 virus and its close coronavirus relatives binds directly to the host angiotensin-converting enzyme 2 (ACE2) to enter host cells. Thus, S protein is a major translational research target, including small molecule inhibitors and extensive vaccine efforts globally (27, 28). Analysis of the gene encoding the S protein identified 41 SNPs, including 25 that produce amino acid changes (**Table 2, Fig. 6A**). Seven of these replacements (A263V, F456L, S967R, T1027I, M1050V, K1157M, and Q1208H) are not represented in the GISAID database as of April 15, 2020. F456 makes contact with the human ACE2 receptor, occupying a pocket formed by ACE2 residues Thr27, Asp30 and Lys31 (**Fig. 6B**) (29). The F456L substitution is expected to detrimentally affect binding to ACE2, as the shorter Leu side chain would not fill the pocket well. We note that seven of the amino acid replacements map to the periphery of the S1 subunit N-terminal domain (NTD). Four of these replacements (A27S, T29I, G261V, and A263V) can be mapped to the recently determined cryo-EM structure of the SARS-CoV-2 spike (27), whereas three other amino acid changes (L18F, T22I, and H69Y) are located in flexible NTD loops that could not be modeled (**Fig. 6A**). The clustering of amino acid replacements to a distinct region of the protein, together with the occurrence of two different amino acid replacements occurring at the same residue is a strong signal of positive selection. Inasmuch as infected patients make antibodies against the NTD region, we favor the idea that host immune selection is a key force contributing to amino acid variation in this region, resulting in an enhanced fitness phenotype of virus variants. Also of note, the D614G replacement was observed in 70% (224 of 320 strains) of the strains sequenced in this study. Residue D614 is located in subdomain 2 (SD-2) of the S protein, and it forms a hydrogen bond and electrostatic interaction with two residues in the S2 subunit of a neighboring protomer (**Fig. 6C**). Replacement of aspartate with glycine would eliminate both interactions, thereby substantively weakening the contact between the S1 and S2 subunits. We speculate that this weakening produces a more fusogenic spike protein, because S1 must first dissociate from S2 before the S2 subunit can refold and mediate fusion of viral and cell membranes. Stated another way, virus strains with the D614G variant may be better able to enter host cells, potentially resulting in enhanced spread.

**Table 2.** Nonsynonymous SNPs in SARS-CoV-2 S protein.

| **Genomic locus** | **S Locus** | **Amino acid change** | **Spike domain<sup>a</sup>** | **Strains (n=320)** |
| --- | --- | --- | --- | --- |
| 21575 | C13T | L5F | S1 | 4 |
| 21614 | C52T | L18F | S1 | 1 |
| 21627 | C65T | T22I | S1 | 2 |
| 21641 | G79T | A27S | S1 | 2 |
| 21648 | C86T | T29I | S1 | 1 |
| 21767 | C205T | H69Y | S1 | 1 |
| 22344 | G782T | G261V | S1 | 1 |
| 22350 | C788T | A263V | S1 | 1 |
| 22928 | T1366C | F456L | S1 / RBD | 1 |
| 23120 | G1558T | A520S | S1 / RBD | 1 |
| 23277 | C1715T | T572I | S1 | 2 |
| 23403 | A1841G | D614G | S1 | 224 |
| 23453 | C1891T | P631S | S1 | 1 |
| 23593 | G2031T | Q677H | S1 | 1 |
| 23755 | G2193T | M731I | S2 | 3 |
| 24077 | G2515T | D839Y | S2 | 1 |
| 24463 | C2901A | S967R | S2 | 1 |
| 24642 | C3080T | T1027I | S2 | 1 |
| 24710 | A3148G | M1050V | S2 | 1 |
| 24794 | G3232T | A1078S | S2 | 3 |
| 25032 | A3470T | K1157M | S2 | 1 |
| 25169 | C3607T | L1203F | S2 | 1 |
| 25186 | G3624T | Q1208H | S2 | 1 |
| 25217 | G3655T | G1219C | S2 | 1 |
| 25350 | C3788T | P1263L | S2 | 1 |

<sup>a</sup>S1 domain; RBD, receptor binding domain; S2 domain

**Fig. 6.**
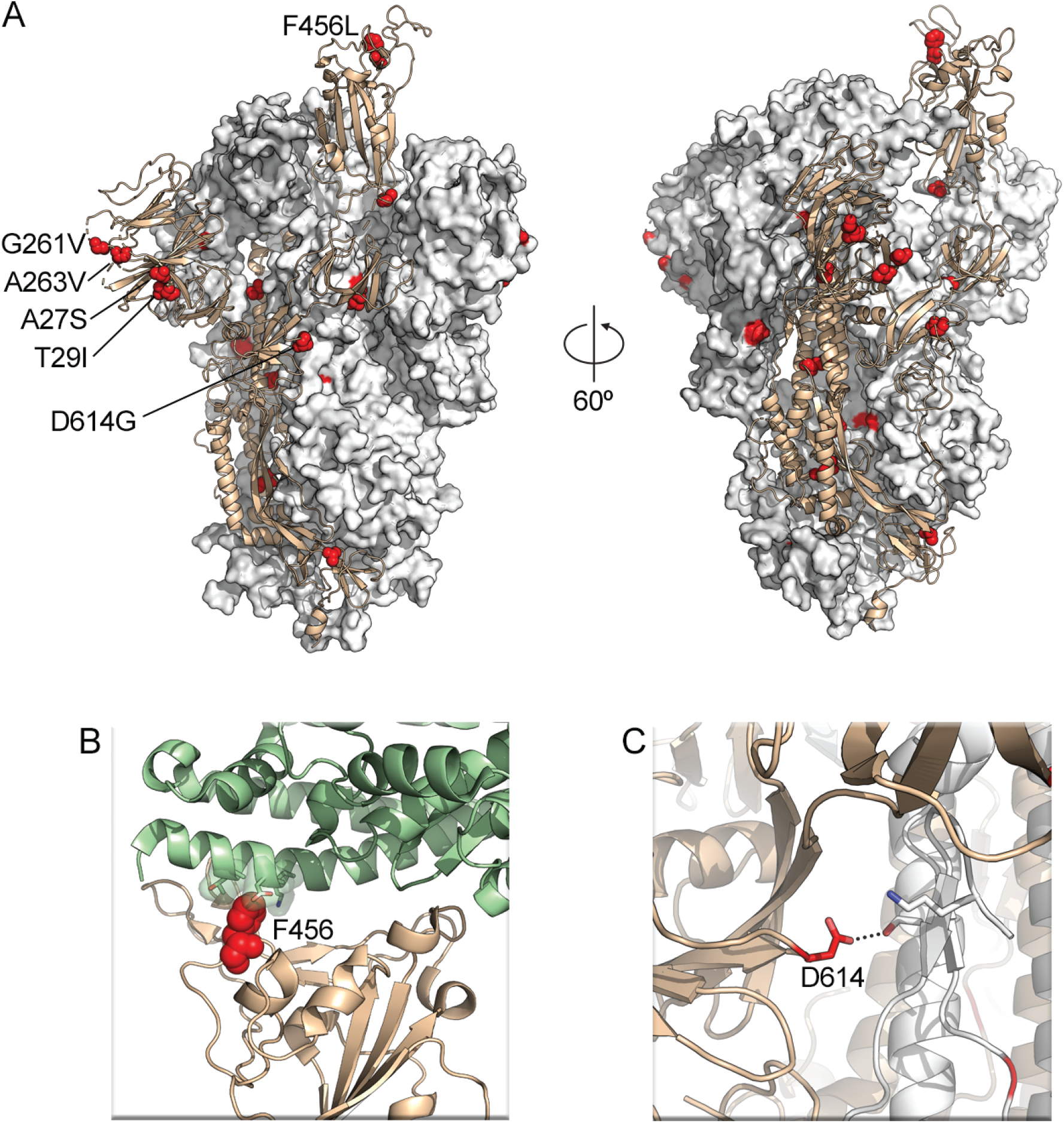
Spike protein amino acid variants. (A) Amino acid replacements identified in this study are shown as red spheres on the structure of the SARS-CoV-2 spike protein. One protomer is shown as ribbons colored tan, whereas the other two protomers are shown as white and gray molecular surfaces. (B) Interface between ACE2 (light green) and spike RBD (tan), shown as ribbons. Spike residue F456 is shown as red spheres. ACE2 residues T27, D30, and K31 are shown as sticks with transparent molecular surfaces. (C) Interaction of D614 with the side chains of two amino acid residues located on the S2 subunit of a different protomer. A hydrogen bond is depicted as a dotted black line.

## Discussion

We discovered substantial genetic diversity of the SARS-CoV-2 viruses causing COVID-19 disease in Houston, Texas. The majority of cases studied are progeny of strains that cause widespread disease in European and Asian countries. We interpret these data as demonstrating that the SARS-CoV-2 virus was introduced into Houston many times independently, likely by individuals who had traveled to different parts of the world. In support of this interpretation, the first cases in metropolitan Houston were associated with a travel history to a known COVID-19 region (14). The data are consistent with the fact that Houston is a large international city characterized by a multi-ethnic population.

The diversity present in our 320 viral genomes contrasts somewhat with data reported recently by Gonzalez-Reiche et al. (30), who studied 84 SARS-CoV-2 isolates causing disease in patients in the New York City region. Those investigators concluded that the vast majority of disease was caused by progeny of strains imported from Europe. Similarly, Bedford et al. (31) reported that much of the COVID-19 disease in the Seattle, Washington area was caused by strains that are progeny of a virus strain recently introduced from China.

The viral diversity present in metropolitan Houston permitted us to test the hypothesis that distinct viral clades were nonrandomly associated with hospitalized COVID-19 patients or disease severity. Although our sample size is relatively small, we found little evidence to support this hypothesis. Clearly, this important matter warrants further study with a larger sample size, and this analysis is currently underway.

We used machine learning classifiers in an attempt to identify SNPs that contribute to increased infection severity or otherwise affect virus-host outcome. The models could not be trained to accurately predict these outcomes from the viral genome sequence data, which may be due to the small sample size and class imbalance. As such, no particular SNPs were identified that are predictive of disease severity or infection outcome. Classifiers were also trained to search for predictors of host characteristics such as age by decade, gender, and ethnic group. We found that the African American versus Caucasian populations had a predictive signal, and hence potentially significant SNPs, which may be borne out with increased sampling in these two groups. However, examination of the geographic distribution of the viral isolates classified using this model largely coincided with the demographic distribution of ethnic groups in the Houston metropolitan region. As such, the underlying SNPs found by the model may reflect social networks present early in the spread of SARS-CoV-2 in the Houston metropolitan area, rather than distinct viral or human genetic factors.

Remdesivir is a nucleoside analog that has been reported to have activity against MERS-CoV, a coronavirus related to SARS-CoV-2 (21). Recently, Grein et al. (22) reported that this drug shows promise in treating COVID-19 patients in a relatively small compassionate use protocol. Because *in vitro* resistance of SARS-CoV to remdesivir has been reported to be caused by either of two amino acid replacements in RdRp (Phe476Leu and Val553Leu), we interrogated our data for polymorphisms in the nsp12 gene. Although we identified 11 different inferred amino acid replacements in the 320 genomes analyzed, none of these were located at the two positions associated with resistance *in vitro*. These findings suggest that if remdesivir proves to be efficacious in COVID-19 patients, and is deployed widely as a treatment, the majority of SARS-CoV-2 strains currently circulating should be susceptible to this drug. However, the Ala448Val polymorphism we identified occurs at an amino acid site that intriguingly is located directly above the nucleotide substrate entry channel and nucleotide binding residues Lys545, Arg553, and Arg555 (20, 25) (**Fig. 5**). One possibility is that substitution of the smaller alanine residue with the bulkier valine may impose structural constraints for the modified nucleotide analog to bind and thereby disfavor remdesivir binding. This in turn leads to reduced incorporation of remdesivir into the nascent RNA, increased fidelity of RNA synthesis, and thus drug resistance. A similar mechanism has been proposed for a V553L change (20). We also identified one strain with a Lys477Asn replacement in nsp12. This substitution is located very close to a Phe476Leu replacement reported to produce partial resistance to remdesivir *in vitro* in SARS-CoV patients from 2004, although the amino acid positions are numbered differently in SARS-CoV and SARS-CoV-2. Structural studies have suggested that this amino acid is surface-exposed, and distant from known key functional elements. Our observed Lys477Asn change is also located in a conserved motif described as a finger domain of RdRp (**Fig. 5**). One speculative possibility is that Lys477 is involved in binding a yet unidentified cofactor such as nsp7 or nsp8, an interaction that could modify nucleotide binding and/or fidelity at a distance. These data warrant additional study in larger patient cohorts, especially individuals treated with remdesivir.

Analysis of the gene encoding the spike protein identified many new inferred amino acid replacements not present in available databases. These data, coupled with structural information available for S protein, raises the possibility that many of the amino acid variants have functional consequences. For example, clustering of several amino acid changes to the NTD suggests varying residues in this region bestow a fitness phenotype. Regardless, we are now beginning to acquire critical information about the location and extent of amino acid replacements occurring in the S protein in natural populations of SARS-CoV-2. These data permitted generation of many biomedically relevant hypotheses now under study.

### Concluding statement

Our work represents analysis of the largest sample of SARS-CoV-2 genome sequences to date from patients in the southern United States. This investigation was facilitated by the fact that we had rapidly assessed the suitability and performance of the SARS-CoV-2 molecular diagnostic test in January 2020, more than a month before the first COVID-19 patient was diagnosed in Houston. Our large healthcare system has eight hospitals and many clinics located in geographically diverse areas of the city. These facilities serve patients of very diverse ethnicities and socioeconomic status, thus our data likely represent a broad overview of virus diversity causing infections throughout metropolitan Houston. We acknowledge that not every “twig” of the SARS-CoV-2 evolutionary tree in Houston is represented in these data because the samples studied were not comprehensive.

The genomes reported here represent the first strains documented to cause COVID-19 disease in the Houston area. Thus, they are an important resource that will underpin further and our ongoing study of SARS-CoV-2 molecular evolution and dissemination in Houston. As of April 15, 2020, there were 5,602 reported cases of COVID-19 in metropolitan Houston, with the number of cases growing daily. We are now sequencing virus genomes essentially in real time from infected patients, which will permit us to provide important data that can be exploited to facilitate an enhanced public health response to this pandemic.

Several regions have reported a flattening of the COVID-19 infection curve, for example the New York City area, China, Italy, and Iceland. This trend suggests that public health and laboratory measures to decrease viral spread are showing promise. In Houston, analogous epidemiologic data have recently been reported. If a decreased virus dissemination rate is successfully maintained, it will be especially important to combine traditional public health control approaches (i.e., case identification and contact tracing) with genome sequence information to deconvolute patterns of infection spread. The availability of extensive viral genome data dating from the earliest reported cases of COVID-19 in metropolitan Houston, coupled with the database we have now constructed, may provide important insight into the origin of new infection spikes occurring as social distancing constraints are relaxed and the region begins to re-open. The genome data will also be useful in assessing ongoing molecular evolution in the spike and other proteins, as herd immunity is generated, either by natural exposure to the SARS-CoV-2 virus or vaccination. The signal of positive selection contributing to diversity in the spike protein is especially concerning and warrants close, intense attention.

## Materials and Methods

### Patient specimens

All specimens were obtained from individuals who were registered patients seen at a Houston Methodist Hospital or associated facilities (e.g., urgent care centers) in the greater metropolitan region. Virtually all individuals met the criteria specified by the Centers for Disease Control and Prevention to be classified as a person under investigation (PUI).

### SARS-CoV-2 molecular testing

Specimens obtained from symptomatic patients with a high degree of suspicion for COVID-19 disease were tested in the Molecular Diagnostics Laboratory at Houston Methodist Hospital using an assay granted Emergency Use Authorization (EUA) from the FDA (https://www.fda.gov/medical-devices/emergency-situations-medical-devices/faqs-diagnostic-testing-sars-cov-2#offeringtests). The assay follows the protocol published by the WHO (https://www.who.int/docs/default-source/coronaviruse/protocol-v2-1.pdf) and uses the 7500 Fast Dx instrument (ABI) and 7500 SDS software (ABI). Testing was performed on material obtained from nasopharyngeal or oropharyngeal swabs immersed in universal transport media (UTM), bronchoalveolar lavage fluid, or sputum treated with dithiothreitol (DTT). To standardize specimen collection, an instructional video was created for Houston Methodist healthcare workers (https://vimeo.com/396996468/2228335d56). Total viral RNA was extracted from patient specimens and tested for the presence of SARS-CoV-2 with an EZ1 virus extraction kit and EZ1 Advanced XL instrument with the Virus card (Qiagen), or QIASymphony DSP Virus kit and QIASymphony instrument (Qiagen).

### SARS-CoV-2 genome sequencing

Libraries for whole viral genome sequencing were prepared according to version 1 of the ARTIC nCoV-2019 sequencing protocol (https://artic.network/ncov-2019). Long reads were generated with the LSK-109 sequencing kit, 24 native barcodes (NBD104 and NBD114 kits), and a GridION instrument (Oxford Nanopore). Short reads were generated with the NexteraXT kit and a MiSeq or NextSeq 550 instrument (Illumina).

### SARS-CoV-2 genome sequence analysis

Consensus viral genome sequences from the Houston isolates were generated using the ARTIC nCoV-2019 bioinformatics pipeline. Publicly available genomes and metadata were acquired through GISAID on April 13, 2020 (16). GISAID sequences containing greater than 1% N characters, and Houston sequences with greater than 5% N characters were removed from consideration. Identical GISAID sequences originating from the same geographic location with the same collection date were also removed from consideration to reduce redundancy. Nucleotide sequence alignments for the combined Houston and GISAID strains were generated using MAFFT version 7.130b with default parameters (32). Sequences were manually curated in JalView (33) to trim the ends and to remove sequences containing spurious inserts. Phylogenetic trees were generated using FastTree with the generalized time-reversible model for nucleotide sequences (34). Trees were parsed, annotated, and visualized using the R packages treeio and ggtree (35–37). CLC Genomics Workbench (QIAGEN) was used to generate the trees in the supplemental figures. Genomes from the Houston strains are available in the EpiCoV collection on GISAID (gisaid.org).

### Analysis of the nsp12 polymerase and S protein genes

The nsp12 viral polymerase and S protein genes were analyzed by plotting SNP density in the consensus alignment using Python (Python v3.4.3, Biopython Package v1.72). The frequency of the SNP in the Houston isolates was assessed, along with the amino acid change for nonsynonymous SNPs.

### Epidemiologic curve

The number of confirmed COVID-19 positive cases was obtained from USAFacts.org (https://usafacts.org/visualizations/coronavirus-covid-19-spread-map/) for Austin, Brazoria, Chambers, Fort Bend, Galveston, Harris, Liberty, Montgomery, and Waller counties. Positive cases for Houston Methodist Hospital patients were obtained from the Laboratory Information System and plotted using the documented collection time.

### Geospatial mapping

The home address zip code for all SARS-CoV-2 positive patients was used to generate a frequency heat map using the Microsoft Power BI Filled map visualization. The home address for patients whose isolates were sequenced was matched to a dictionary of addresses downloaded from OpenAddresses.io (https://openaddresses.io/) to obtain the latitude and longitude geocoding data. Because addresses are transcribed and subject to manual error, the fuzzywuzzyR package was used to match the best address in the dictionary. All address matches were manually reviewed for accuracy. The latitude/longitude coordinates were plotted onto a map using the Microsoft Power BI Desktop Map visualization.

To examine geographic relatedness among genetically similar isolates, the geospatial maps were filtered to four small independent clades. The home address of the patients was again visualized using the Microsoft Power BI Desktop Map visualization.

### Time series

The geospatial data were filtered into three time intervals (3/5/2020-3/16/2020, 3/5/2020-3/23/2020, and 3/5/2020-3/30/2020) to illustrate the spread of confirmed SARS-CoV-2 positive patients we identified over time.

### Machine learning analysis

Machine learning models were trained to predict patient metadata categories including mortality, length of stay, inpatient versus outpatient status, ICU admission, overall outcome, gender, age, and ethnicity from viral sequence data. Models were trained with the consensus whole genome FASTA files by dividing each viral genome into 15-mer oligonucleotide k-mers, which were used as features to train XGBoost models (38) as described previously (39, 40). Models were assessed by computing F1 scores for classifiers and R^2^ scores for regression models. Unless otherwise stated, data for the first 5-folds of a 10-fold cross validation are shown.

## ACKNOWLEDGMENTS

We thank Dr. Steven Hinrichs and colleagues at the Nebraska Public Health Laboratory, and Dr. David Persse and colleagues at the Houston Health Department for providing samples used to validate our SARS-CoV-2 virus molecular assay. We thank Drs. Jessica Thomas and Zejuan Li, Erika Walker, the very talented and dedicated molecular technologists, and the many labor pool volunteers in the Molecular Diagnostics Laboratory for their dedication to patient care. We also thank Brandi Robinson, Harrold Cano, and Cory Romero for technical assistance. We are indebted to Drs. Marc Boom and Dirk Sostman for their support, and to many very generous Houston philanthropists for their tremendous support of this ongoing project, including but not limited to anonymous, Ann and John Bookout III, Carolyn and John Bookout, Ting Tsung and Wei Fong Chao Foundation, Ann and Leslie Doggett, Freeport LNG, the Hearst Foundations, Jerold B. Katz Foundation, C. James and Carole Walter Looke, Diane and David Modesett, the Sherman Foundation, and Paula and Joseph C. “Rusty” Walter III. We gratefully acknowledge the originating and submitting laboratories of the SARS-CoV-2 genome sequences from GISAID’s EpiFlu™ Database used in some of the work presented here. We also thank many colleagues for critical reading of the manuscript and suggesting improvements, and Sasha Pejerrey, Adrienne Winston, Heather McConnell, and Kathryn Stockbauer for editorial contributions. We are especially indebted to Drs. Nancy Jenkins and Neal Copeland for their scholarly suggestions to improve the manuscript.

## Author contributions

J.M.M. designed the project; S.W.L, R.J.O., P.A.C. D.W.B., J.J.J., M.S., M.N., M.O.S., C.C.C., P.Y., L.P., S.S., H.H., G.E., M.K., and J.S.M. performed research. All authors contributed to writing the manuscript.

This study was supported by the National Institutes of Health grants AI146771-01 and AI139369-01, and the Fondren Foundation, Houston Methodist Hospital and Research Institute (to JMM), and NIH grant AI127521 (to J.S.M.). J.J.D, M.S., and M.N. are supported by the NIAID Bacterial and Viral Bioinformatics resource center award (contract number 75N93019C00076). JJD and MN are also funded by the United States Defense Advanced Research Projects Agency Award iSENTRY Friend or Foe program award (contract number HR0011937807).

## Supplemental Figures

**Supplemental Fig. 1-7. Cladograms labeled with patient data, including (age, S1; gender, S2, ethnicity, S3, level of care, S4; length of stay, S5; days on ventilator, S6; and mortality, S7)**.

**Supplemental Fig. 8**. SARS-CoV-2 strains with a Met600Ile polymorphism in nsp12 are located in a distinct genetic cluster. The 29 strains containing the Met600Ile polymorphism are highlighted in red on the cladogram between 2 o’clock and 3 o’clock.

**Supplemental Table 1.**
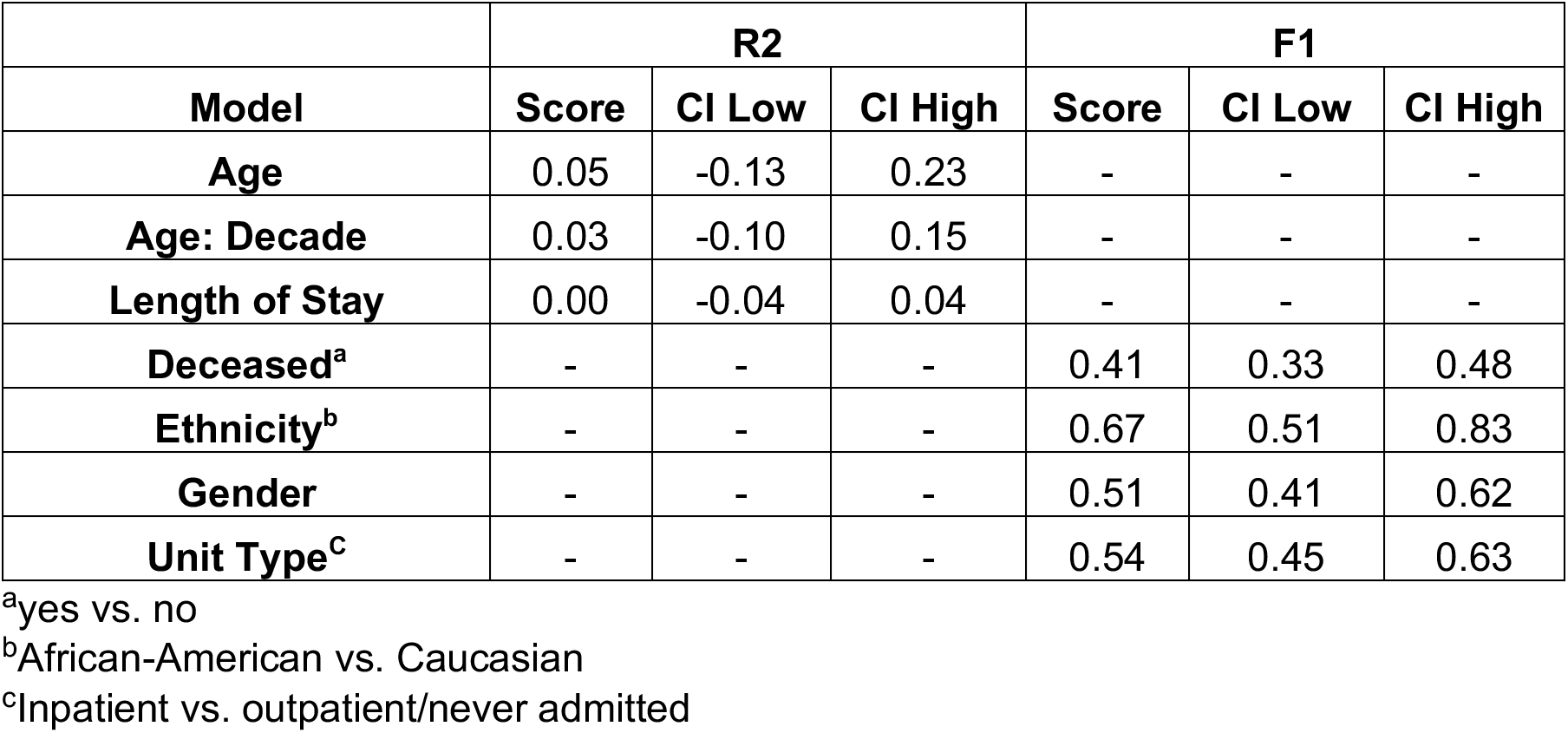
Classifier accuracy scores and performance information.

| **Model** | **R2** |  |  | **F1** |  |  |
| --- | --- | --- | --- | --- | --- | --- |
|  | **Score** | **CI Low** | **CI High** | **Score** | **CI Low** | **CI High** |
| **Age** | 0.05 | -0.13 | 0.23 | - | - | - |
| **Age: Decade** | 0.03 | -0.10 | 0.15 | - | - | - |
| **Length of Stay** | 0.00 | -0.04 | 0.04 | - | - | - |
| **Deceased<sup>a</sup>** | - | - | - | 0.41 | 0.33 | 0.48 |
| **Ethnicity<sup>b</sup>** | - | - | - | 0.67 | 0.51 | 0.83 |
| **Gender** | - | - | - | 0.51 | 0.41 | 0.62 |
| **Unit Type<sup>c</sup>** | - | - | - | 0.54 | 0.45 | 0.63 |

<sup>a</sup>yes vs. no
<sup>b</sup>African-American vs. Caucasian
<sup>c</sup>Inpatient vs. outpatient/never admitted

