## Supplemental Figures for "Molecular Architecture of Early Dissemination and Evolution of the SARS-CoV-2 Virus in Metropolitan Houston, Texas"

Supplemental Figure 1

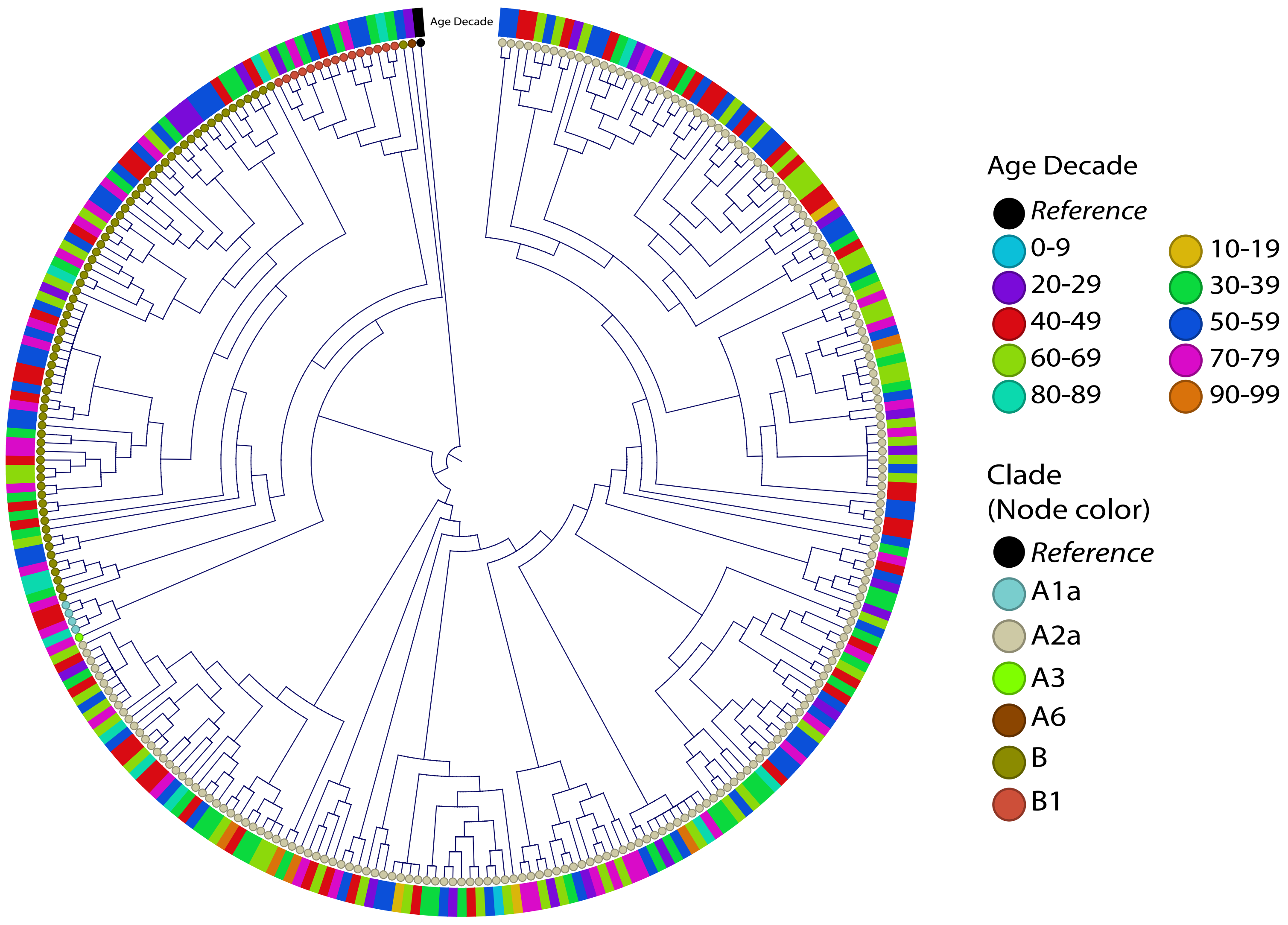

Supplemental Figure 2

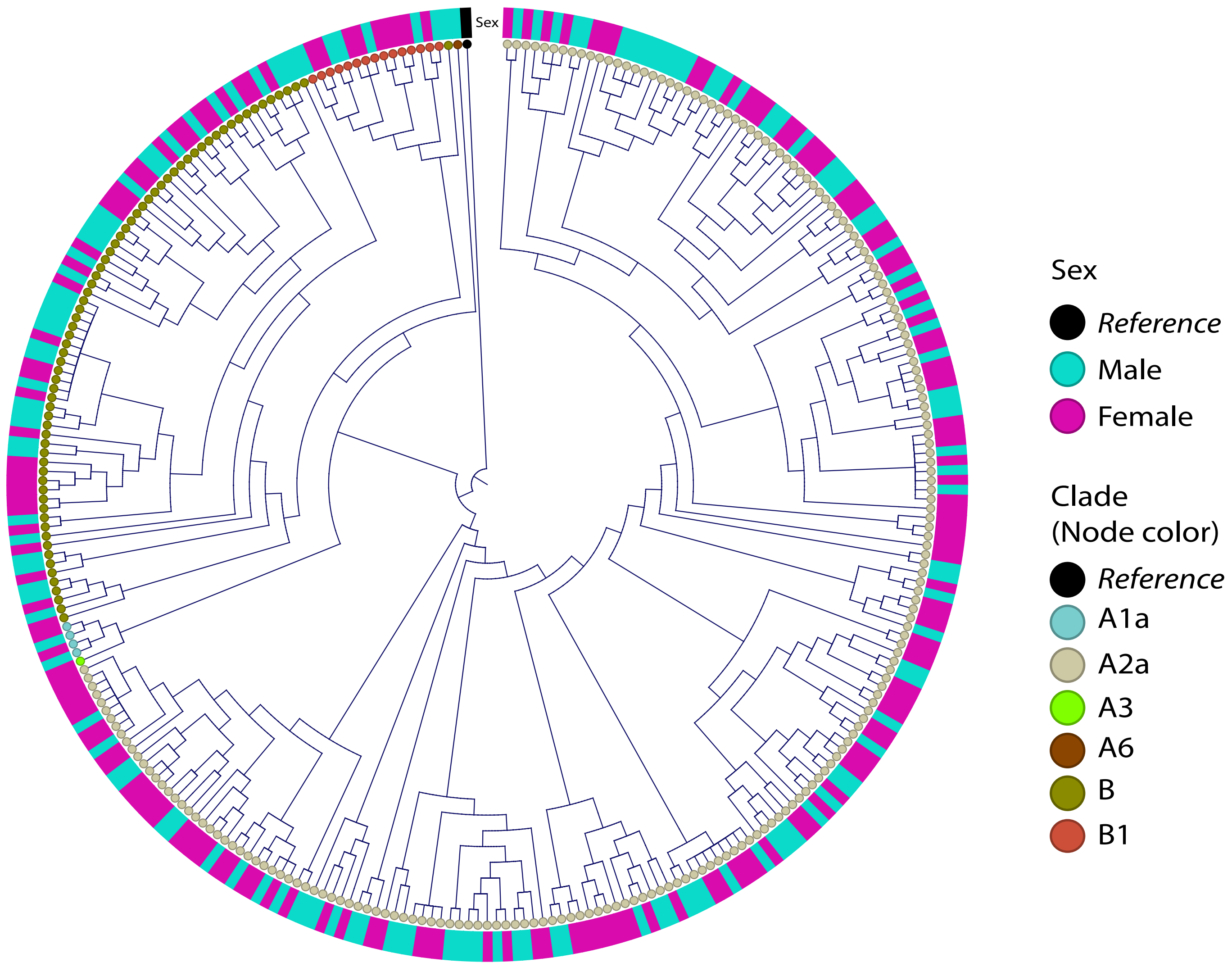

Supplemental Figure 3

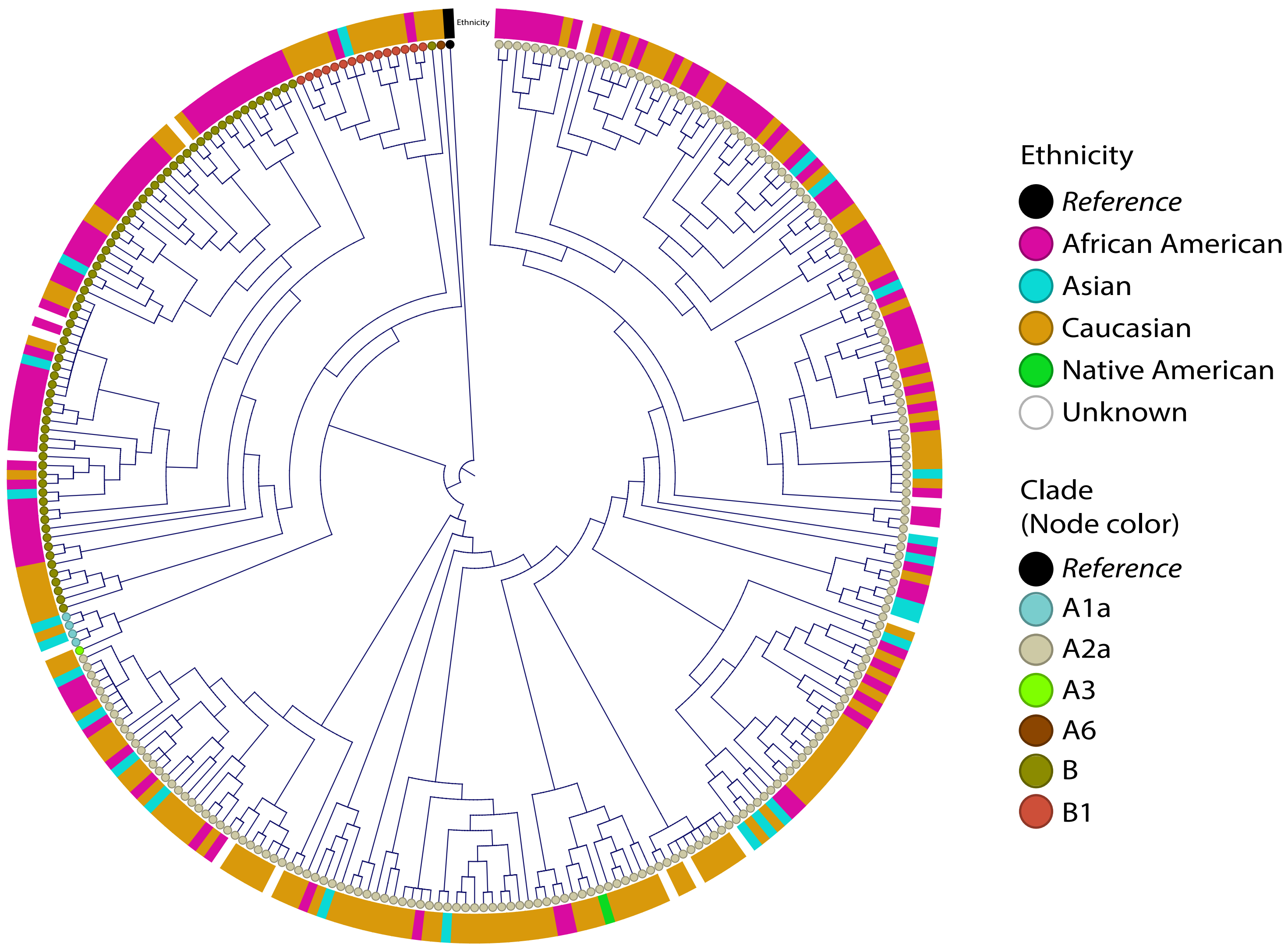

Supplemental Figure 4

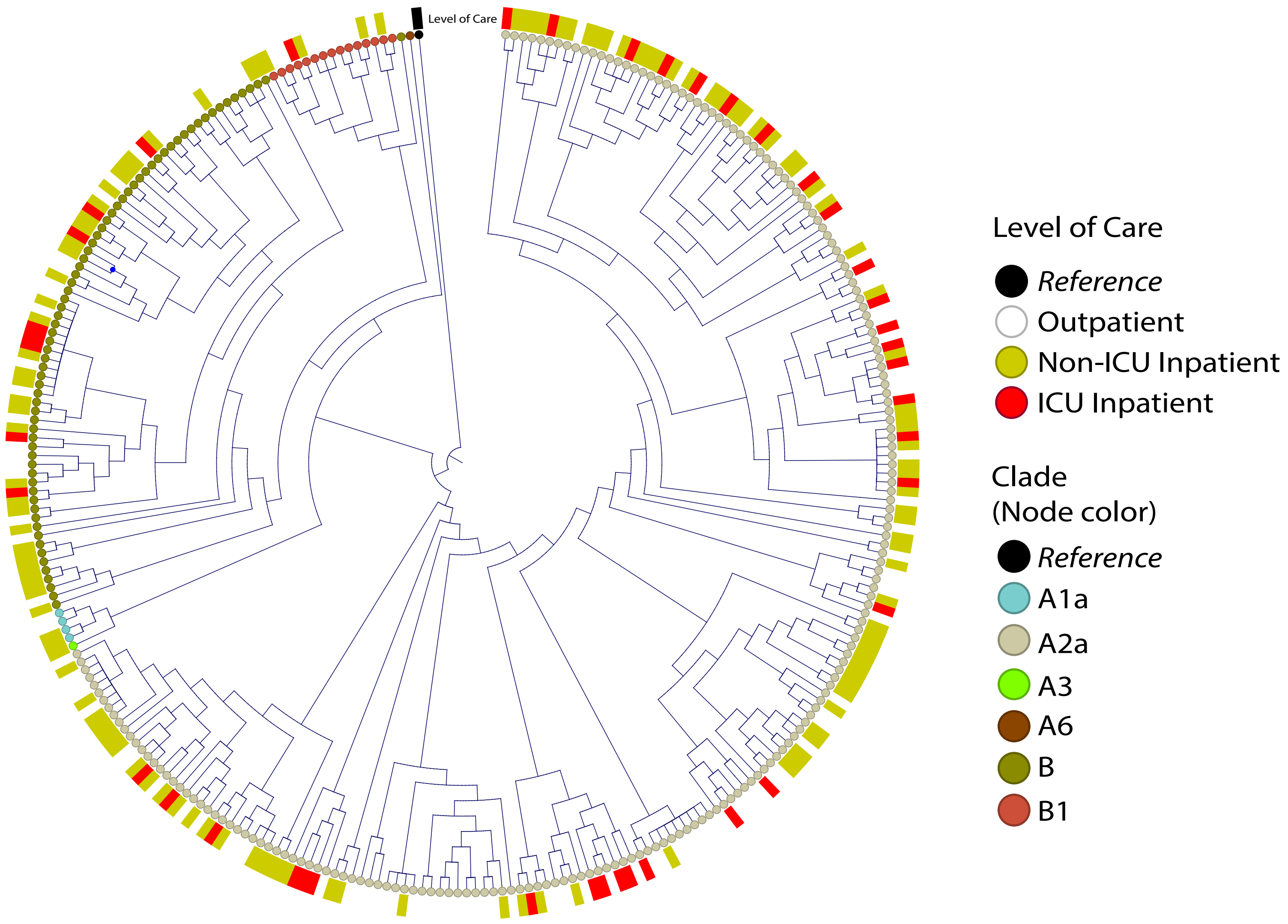

Supplemental Figure 5

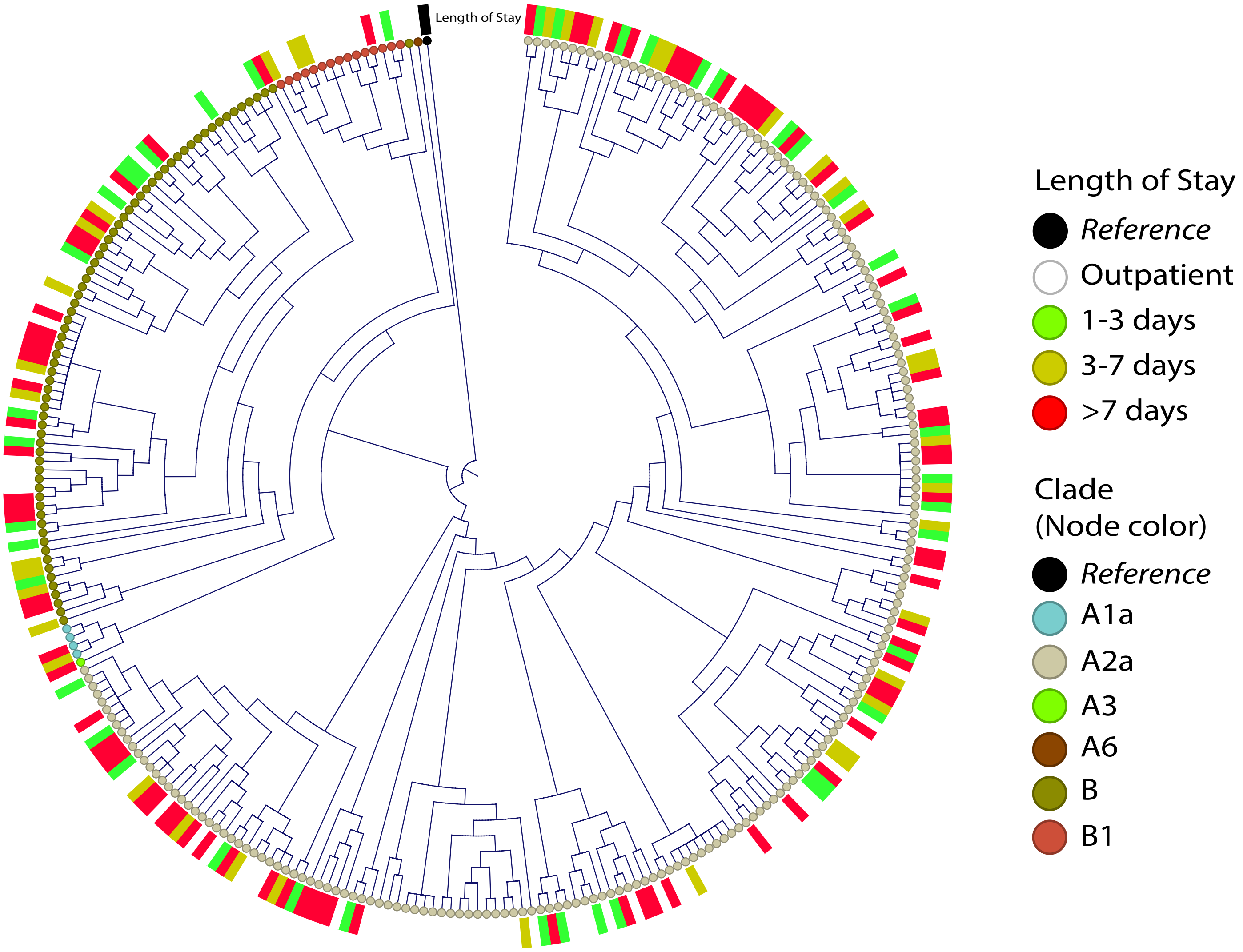

Supplemental Figure 6

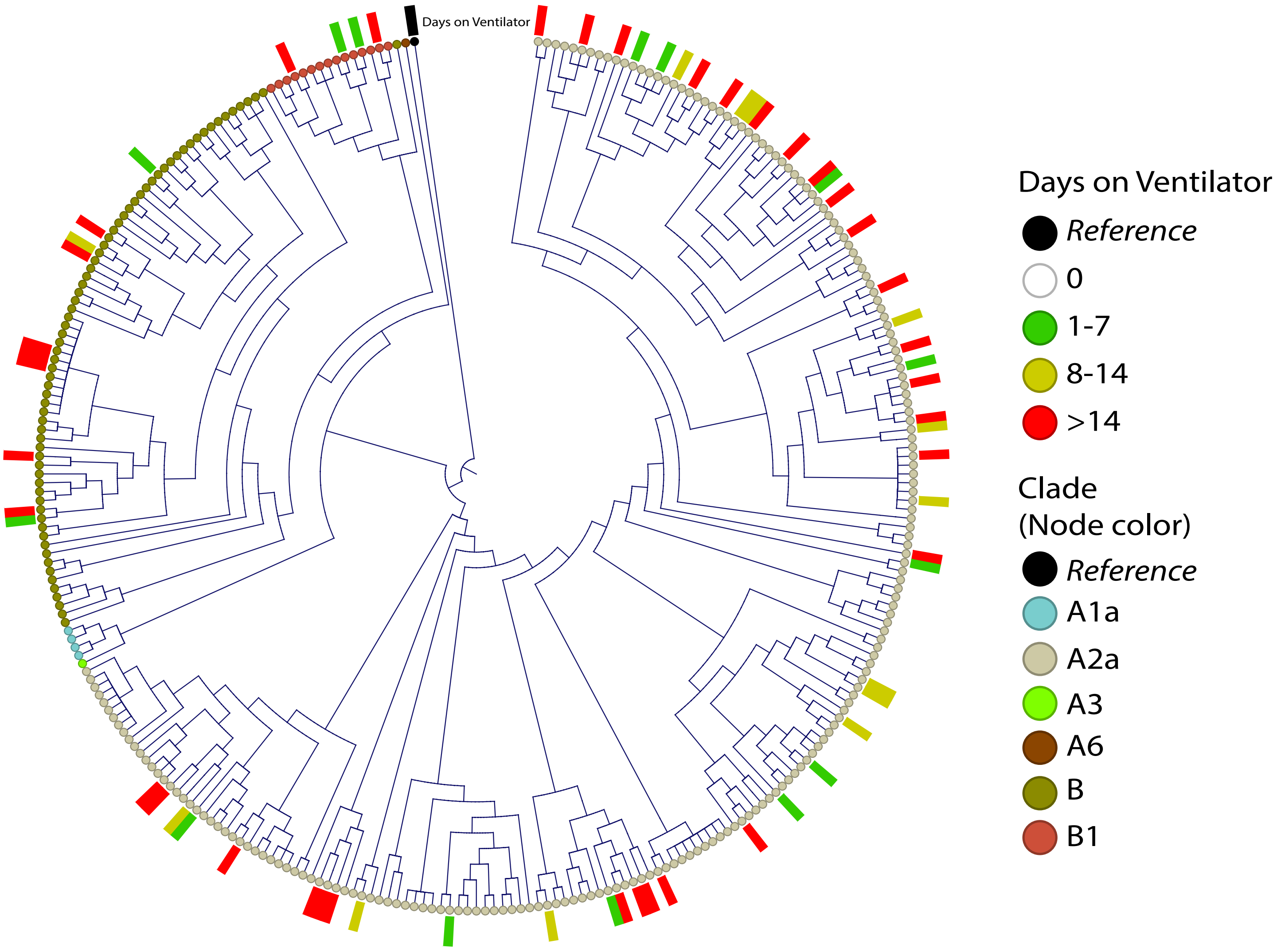

Supplemental Figure 7

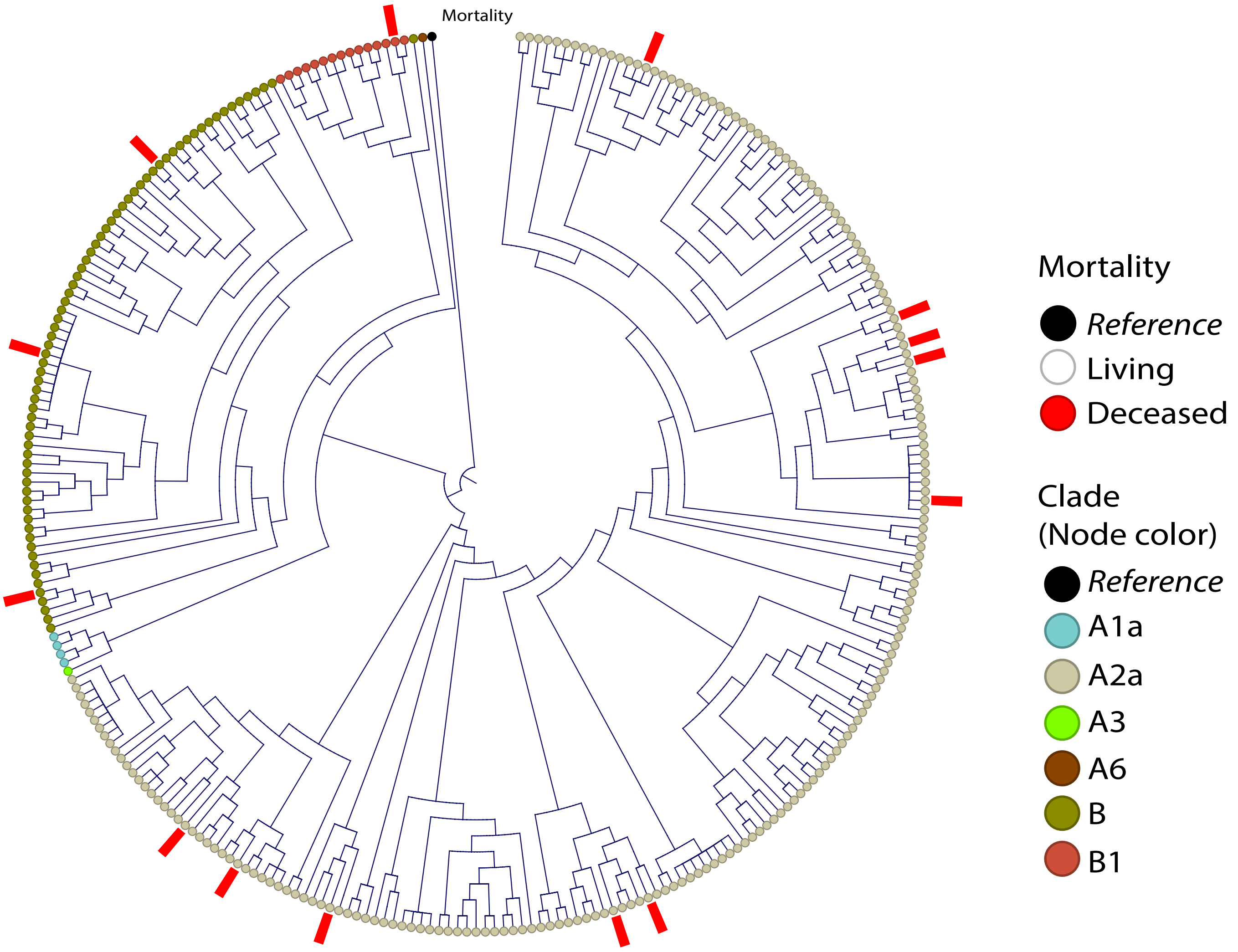

Supplemental Figure 8

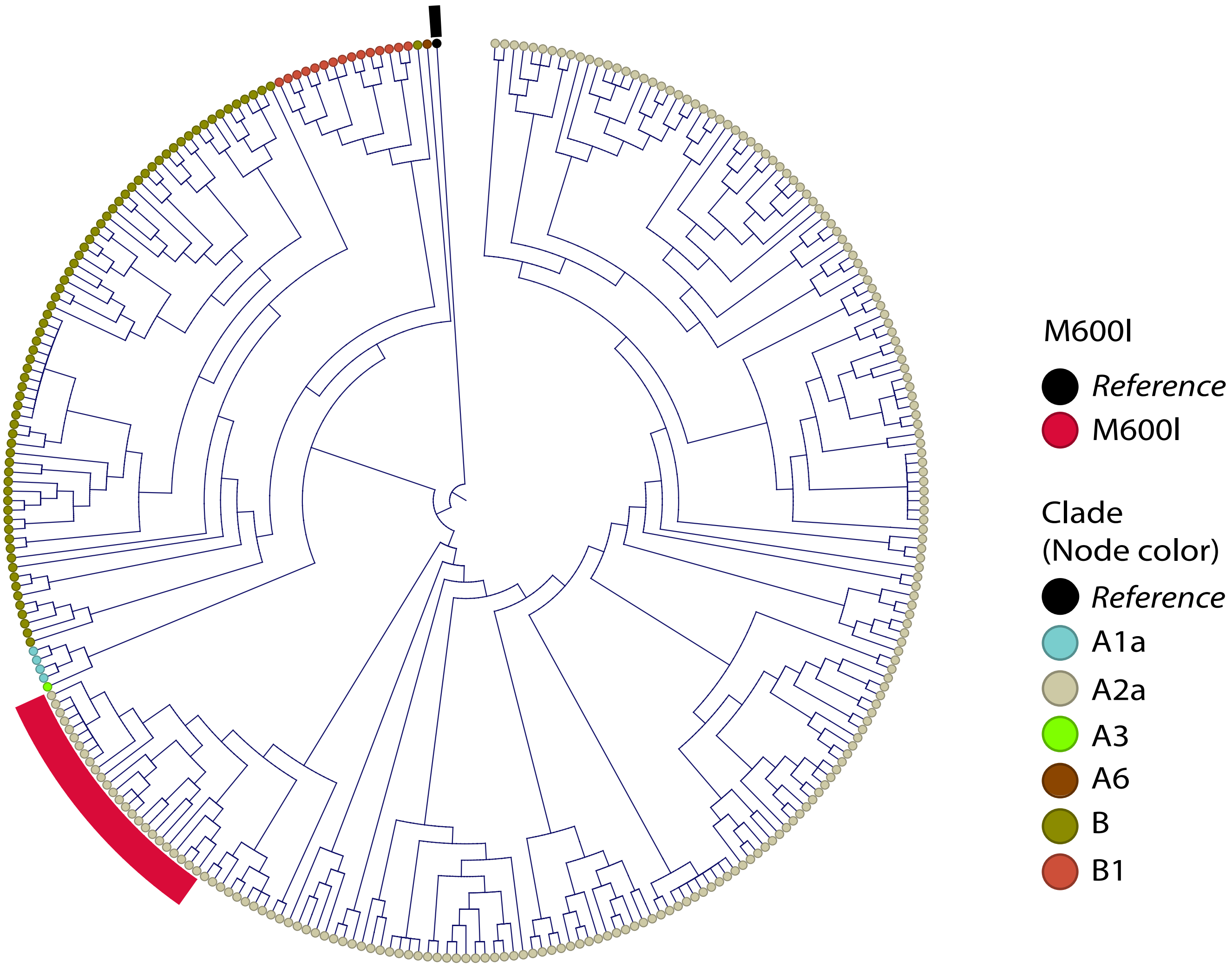
